# Negative cooperativity underlies dynamic assembly of the Par complex regulators Cdc42 and Par-3

**DOI:** 10.1101/2022.08.12.503756

**Authors:** Elizabeth Vargas, Kenneth E. Prehoda

**Affiliations:** Institute of Molecular Biology, Department of Chemistry and Biochemistry, 1229 University of Oregon, Eugene, OR 97403

## Abstract

The Par complex polarizes diverse animal cells through the concerted action of multiple regulators. Binding to Par-3 couples the complex to cortical flows that construct the Par membrane domain. Once localized properly, the complex is thought to transition from Par-3 to the Rho GTPase Cdc42 to activate the complex. While this transition is a critical step in Par-mediated polarity, little is known about how it occurs. Through a biochemical reconstitution approach utilizing purified, intact Par complex and qualitative binding assays, we found that Par-3 and Cdc42 exhibit strong negative cooperativity for the Par complex. We show that the interactions between the second and third PDZ protein interaction domains of Par-3 and the aPKC Kinase-PBM (PDZ binding motif) mediate the displacement of Cdc42 from the Par complex. Our results indicate that Par-3, Cdc42, Par-6, and aPKC are the minimal components that are sufficient for this transition to occur and that no external factors are required. Our findings provide the mechanistic framework for understanding a critical step in the regulation of Par complex polarization and activity.

## Introduction

The polarization of animal cells by the Par complex is a highly dynamic, multi-step process, that begins when actomyosin-generated cortical flows transport membrane-bound Par complex from cellular regions where it is catalytically inactive to a single cortical domain where it becomes activated (1–8). The transition from an inactive to active complex is mediated by the formation of two distinct complexes: one bound to the multi-PDZ protein Par-3 (Bazooka; Baz in *Drosophila*) and a Rho GTPase Cdc42-bound complex. Par-3 has many reported interactions with both Par complex components, atypical Protein Kinase C (aPKC) and Par-6, whereas Cdc42 has one well-defined binding site on Par-6 (Figure 1A) (9–19). The transition between these two regulators precisely controls Par complex polarization and activity, with Par-3 coupling the Par complex to cortical flow while inhibiting aPKC activity and GTP-bound Cdc42 maintaining the Par complex at the cell cortex while stimulating aPKC activity (5, 20, 21). Despite the critical importance of the transition from Par-3 to Cdc42 in the mechanism of Par-mediated polarity, very little is known about how it occurs.

**Figure 1.**
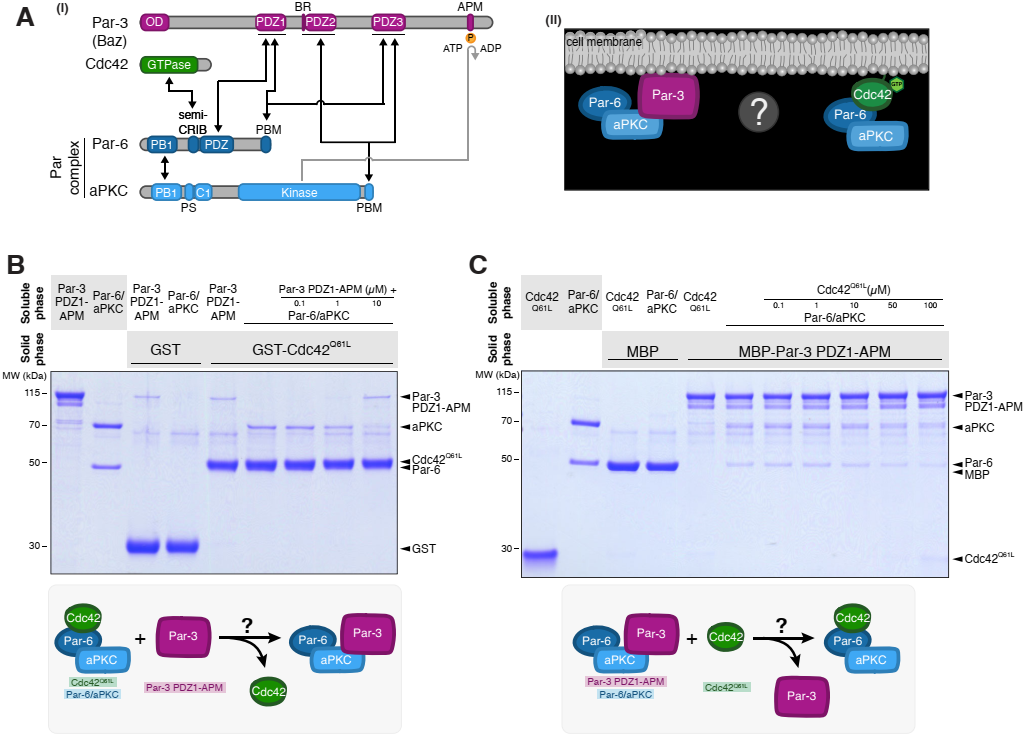
Par-3 and Cdc42 bind to the Par complex with strong negative cooperativity. (**A**) (i) Domain architecture of Par-3, Cdc42, and the Par complex with reported interactions between Par-3, Cdc42, and the Par complex. Black arrows indicate reported interactions, and the grey arrow indicates phosphorylation. (ii) Schematic for the Par complex transition from Par-3 to Cdc42. (**B**) Effect of Par-3 PDZ1-APM on the interaction between Cdc42 and the Par complex. Solid-phase (glutathione resin)-bound glutathione S-transferase (GST)-fused Cdc42^Q61L^ (constitutively active Cdc42) incubated with Par complex and/or increasing concentrations of Par-3 PDZ1-APM. Shaded regions indicate the fraction applied to the gel (soluble-phase or solid-phase components after mixing with soluble-phase components and washing). (**C**) Effect of Cdc42 on the interaction between Par-3 and the Par complex. Solid-phase (amylose resin)-bound maltose-bound protein (MBP)-fused Par-3 PDZ1-APM incubated with Par complex and/or increasing concentrations of Cdc42^Q61L^. Labeling as described in (B).

While *in vivo* evidence indicates the Par complex switches from Par-3 to Cdc42-bound states, biochemical evidence suggests that Par-3 and Cdc42 can bind the Par complex simultaneously to form a quaternary complex. A co-immunoprecipitation experiment using cell extracts found that Cdc42-bound Par complexes also contain Par-3 (9). However, *in vivo* evidence indicate that there are two distinct cortical pools of Par complex, colocalizing with either Par-3 or Cdc42, and loss of Cdc42 increases the amount of Par-3-bound complex (3, 5, 20, 22). The reported ability of Cdc42 and Par-3 to bind simultaneously to the Par complex has influenced models for how the transition between the regulators could occur *in vivo*. In one model, Cdc42 briefly docks onto Par-3-bound Par complex and activates aPKC, resulting in the phosphorylation and release of Par-3 from the complex (23–25). However, recent studies show that phosphorylation of Par-3 by aPKC does not dissociate Par-3 from the Par complex (18, 26). In another proposed model, actomyosin contractility mechanically dissociates Par-3 clusters and facilitates the Par complex transition to Cdc42 (5, 20, 21).

Because the available biochemical data suggests that Par-3 and Cdc42 can bind simultaneously to the Par complex, models for the transition between the two regulators necessarily include other mechanisms (e.g. phosphorylation) or cellular components (e.g. actomyosin contractility). However, the limited *in vitro* evidence is based on results from cell extracts or experiments using truncated proteins. Additionally, the numerous reported interactions between Par-3 and the Par complex have made it challenging to understand how the Par-3-bound Par complex is regulated. Finally, very little structural information is known about the Par complex and whether the Par-3 and Cdc42 binding sites are in close proximity to one another to regulate the formation of these complexes. Here we have used a biochemical reconstitution approach with purified components to determine the elements sufficient for Par complex switching between Par-3 and Cdc42. The results provide the mechanistic framework for understanding how the Par complex transitions from Par-3 to Cdc42 to form two distinct complexes.

## Results

### Par-3 and Cdc42 bind with negative cooperativity to the Par complex

Although Par-3 and Cdc42 are thought to form mutually exclusive complexes with the Par complex *in vivo*, they have been shown to bind simultaneously in a co-immunoprecipitation experiment using cell extracts (9). We examined whether Par-3 and Cdc42 influence one another’s binding to the Par complex using a reconstitution system. We performed a qualitative affinity chromatography (pull-down) assay with purified Par complex and Par-3 PDZ1-APM (a fragment containing all known interaction motifs between Par-3 and Par-6/aPKC) and GST-fused Cdc42^Q61L^ (constitutively active). The binding buffer included ATP to ensure that the aPKC kinase domain did not form a stalled complex with its phosphorylation site on Par-3.

We formed a complex of Cdc42-bound Par-6/aPKC by placing GST-Cdc42^Q61L^ on the solid phase and incubating with soluble, purified Par complex. We assessed the effect of Par-3 on the Cdc42-bound Par complex by adding increasing concentrations of Par-3 PDZ1-APM. If Par-3 binding to the Par complex had no effect on Cdc42 binding, or the proteins bound with positive cooperativity, we expected that Par-3 would become part of the solid phase complex and the amount of Par complex adhered to the solid phase would stay the same or increase. Alternately, if the Cdc42 and Par-3 binding sites exhibited negative cooperativity, either via direct steric occlusion or an allosteric mechanism, little or no Par-3 would be part of the Cdc42-bound solid phase complex, and the amount of Par complex on the solid phase would decrease (as the affinity of the Par complex for Cdc42 was reduced by binding to Par-3). We observed that addition of Par-3 significantly reduced the amount of Par complex associated with solid phase Cdc42^Q61L^ (Figure 1B). Furthermore, little or no additional Par-3 appeared in the solid phase relative to a GST control. Our results indicate that in the context of these four proteins, Cdc42 and Par-3 bind with negative cooperativity to Par-6/aPKC.

In a system with two distinct binding sites coupled to one another via negative cooperativity, each protein should reduce the affinity of the Par complex for the other. However, the effect of Cdc42 on Par-3 binding to the Par complex is complicated by the many potential Par-3 binding sites on the Par complex (Figure 1A). In principle, not all Par-3 binding sites could be coupled to Cdc42 binding, a scenario in which addition of Cdc42 to solid phase Par-3-bound Par complex might not significantly alter the amount of solid phase Par complex. To determine if Cdc42 influences Par-3-bound Par complex, we adsorbed Par complex bound to MBP-Par-3 PDZ1-APM to the solid phase and examined the effect of increasing concentrations of Cdc42^Q61L^. We observed that addition of Cdc42 reduced the amount of Par complex associated with solid phase Par-3 and Cdc42 was not significantly incorporated into the solid phase (Figure 1C). Displacement of Par complex from Par-3 required a significantly higher concentration of Cdc42 than we observed for Par-3 displacement of Cdc42-bound complex.

Our results indicate that Par-3 and Cdc42 compete for binding to the Par complex (i.e. negative cooperativity) and that a quaternary complex does not form at levels detectable in our assay. Our results may differ from previous studies using cell extracts because aPKC’s kinase domain is known to form stalled complexes with substrates like Par-3 when ATP is not available to complete the catalytic cycle (forming a persistent interaction rather than a transient interaction) (18, 26). Additionally, other cellular factors could potentially allow Par-3 and Cdc42 to bind to the Par complex simultaneously. In terms of understanding how the Par complex might transition from Par-3 to Cdc42, our results demonstrate that no other proteins are required–Par-3 and Cdc42 alone are sufficient to form mutually exclusive complexes with the Par complex.

### Par-3 binds the Par complex with higher affinity than Cdc42

To understand why Par-3 is more effective at displacing Cdc42 from the Par complex, we measured the binding affinities of Par-3 and Cdc42 for the Par complex using a supernatant depletion assay (27). We recently reported the affinity of Par-3 PDZ1-APM for the Par complex (19), and the affinity of Cdc42 for the Par complex has not been determined. One explanation for the high concentration of Cdc42 required to displace Par-3 from the Par complex is that Cdc42 has a lower affinity for the Par complex compared to Par-3. Utilizing a supernatant depletion assay, we found that Par-3 PDZ1-APM bound with high affinity to the Par complex (K_d_ of 0.62 μM or ΔG° of 8.32 kcal/mol), similar to what we reported previously (Figure 2) (19). We measured a much weaker affinity of Cdc42 for the Par complex (K_d_ of 5.37 μM or ΔG° of 7.08 kcal/mol) (Figure 2). An affinity of 0.05 μM has been reported between Cdc42 and the Par complex, although only for the Par-6 CRIB-PDZ fragment (13). Based on our measurements, Cdc42 binds with ~1.2 kcal/mole lower affinity than Par-3 to the Par complex explaining why Par-3 displaces Cdc42 more efficiently.

**Figure 2.**
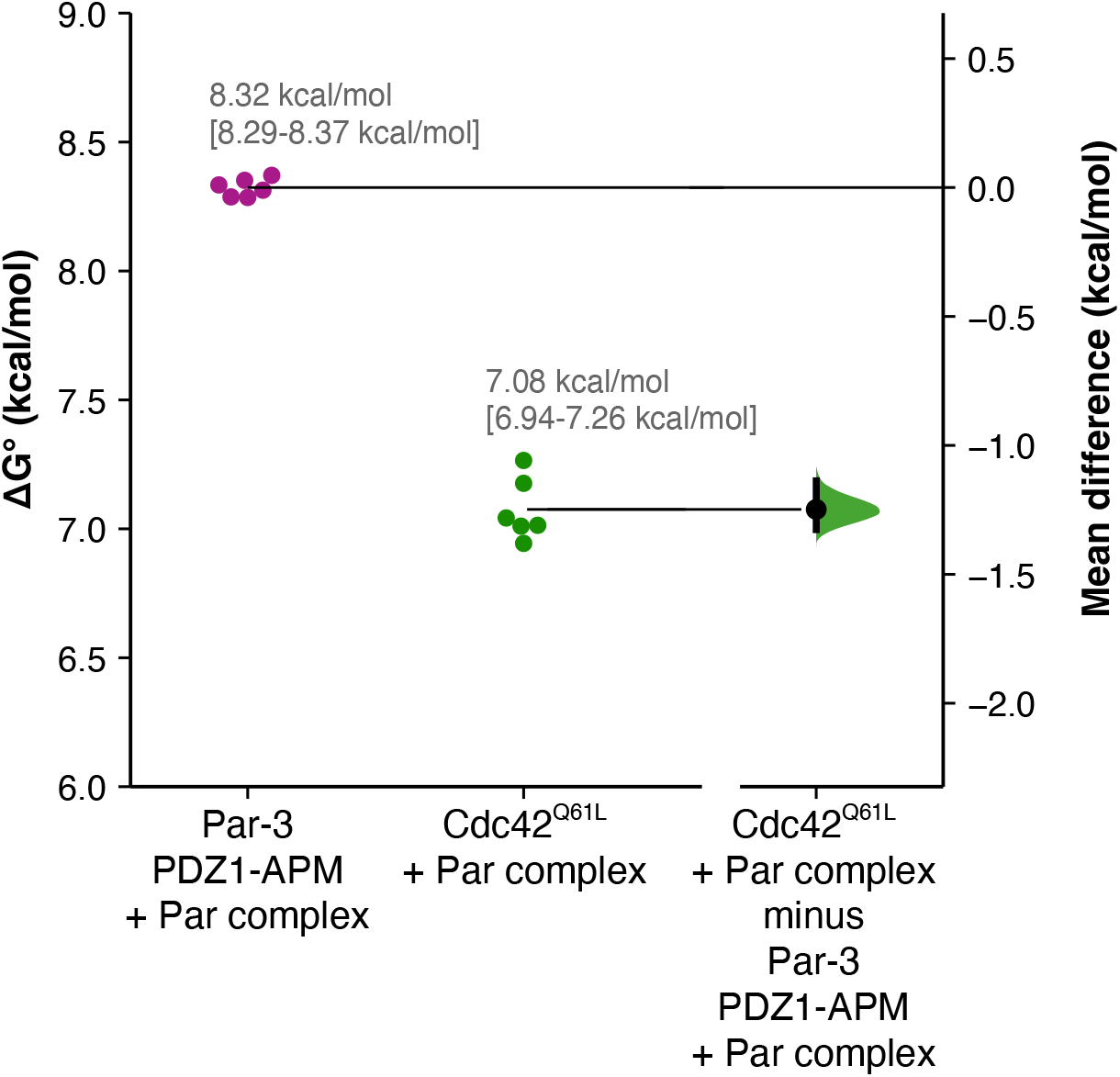
Par-3 binds to the Par complex with a greater affinity than Cdc42. Gardner-Altman estimation plot of Par-3 PDZ1-APM/Par complex and Cdc42^Q61L^/Par complex binding affinities measured using a supernatant depletion assay. The results of each replicate (filled circles) are plotted on the left and the mean difference is plotted on the right as a bootstrap sampling distribution (shaded region) with a 95% confidence interval (black error bar).

### The Par-3 PDZ2—aPKC Kinase-PBM interaction mediates the displacement of Cdc42 from the Par complex

Given that Par-3 and Cdc42 bind with negative cooperativity to Par-6/aPKC, we sought to identify the binding sites on the Par complex that are coupled. While the interaction between Cdc42 and Par-6 semi-CRIB is well established, several interactions between Par-3 and the Par complex have been identified (Figure 1A) (9–19). We excluded the interaction of the aPKC kinase domain with its phosphorylation site on Par-3 (i.e. the Par-3 APM) because the interaction is transient in the presence of ATP, as expected for an enzyme-substrate interaction (18, 26). Given that each Par-3 PDZ domain reportedly interacts with either Par-6 or aPKC, more than one interaction between Par-3 and the Par complex could be involved in displacement of Cdc42 from the Par complex. However, if only one of the interactions between Par-3 and the Par complex displaces Cdc42 from the Par complex, deletion of the required Par-3 element would eliminate Par-3’s negative cooperativity with Cdc42 for the Par complex. Alternately, removal of more than one Par-3 element might be necessary to eliminate displacement of Cdc42 from the Par complex by Par-3. To distinguish between these possibilities, we generated deletions of individual Par-3 PDZ domains in the context of the PDZ1-APM fragment and tested which Par-3 elements are involved in displacing Cdc42 from the Par complex. We examined the effect of Par-3 PDZ1-APM, ΔPDZ1, ΔPDZ2, or ΔPDZ3 on Cdc42-bound Par complex. We did not detect an effect of removing PDZ1 or PDZ3 on Par-3’s ability to displace Cdc42 from the Par complex (Figure 3A). In contrast, deletion of Par-3 PDZ2 eliminated displacement of Cdc42 such that the amount of Par complex associated with solid phase Cdc42 did not change upon addition of Par-3 PDZ1-APM ΔPDZ2 (Figure 3A). Our results indicate that neither Par-3 PDZ1 or PDZ3 are required for negative cooperativity with Cdc42 for the Par complex and that displacement of Cdc42 from the Par complex by Par-3 is dependent on the PDZ2 domain.

**Figure 3.**
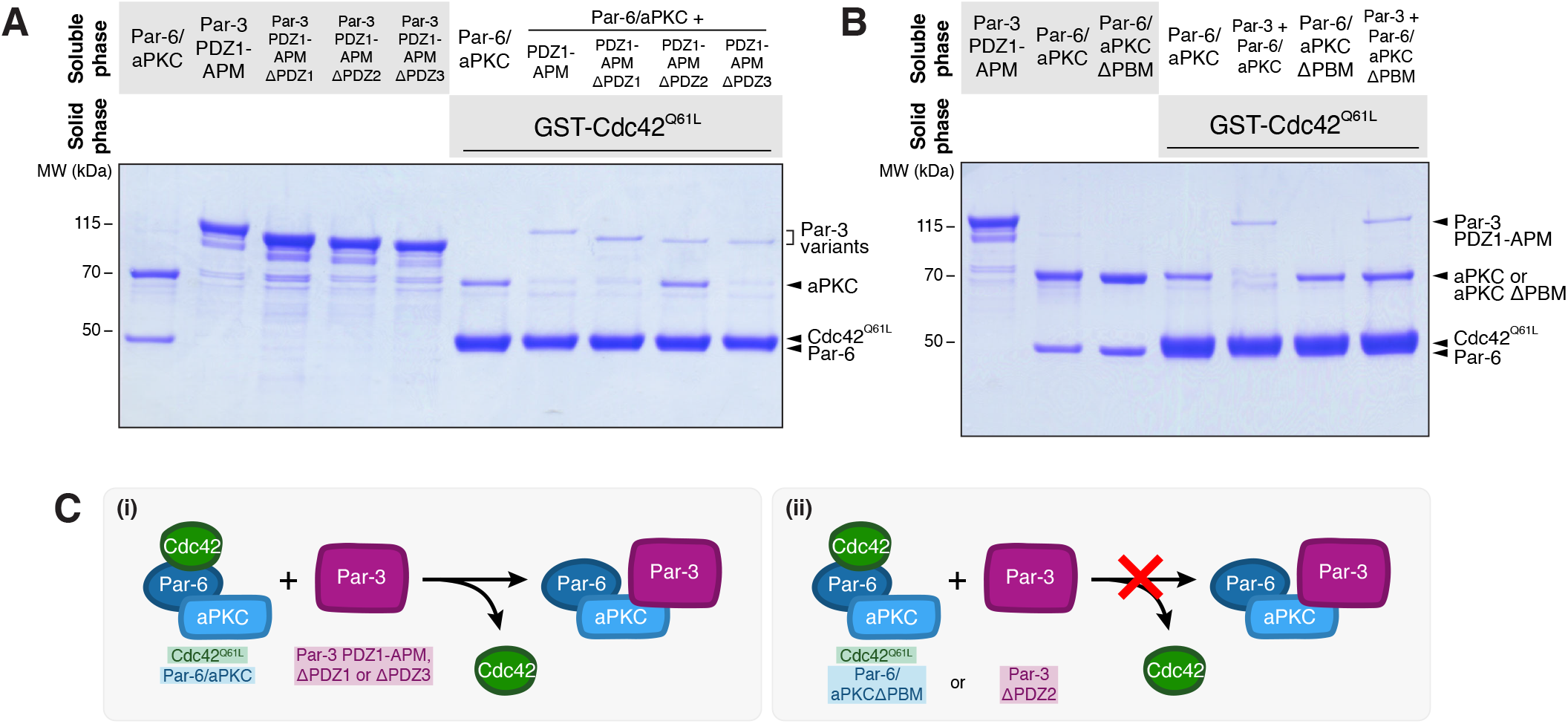
The Par-3 PDZ2 and aPKC PBM interaction is required for facilitating the displacement of Cdc42 from the Par complex. (**A**) Effect of removing individual Par-3 PDZ domains in the context of PDZ1-APM on the displacement of Cdc42 from the Par complex. Solid-phase (glutathione resin)-bound glutathione S-transferase (GST)-fused Cdc42^Q61L^ incubated with Par complex and/or Par-3 PDZ1-APM, ΔPDZ1, ΔPDZ2, or ΔPDZ3. Shaded regions indicate the fraction applied to the gel (soluble-phase or solid-phase components after mixing with soluble-phase components and washing). (**B**) Effect of removing the aPKC PBM in the context of the intact Par complex on Par-3’s ability to displace Cdc42 from the Par complex. GST-fused Cdc42^Q61L^ incubated with Par complex or Par complex ΔaPKC PBM and/or Par-3 PDZ1-APM. Labeling as described in (A). (**C**) Summary of Cdc42, Par-3, Par-6/aPKC elements that are necessary for negative cooperativity between Par-3 and Cdc42 for the Par complex.

We recently discovered that Par-3 PDZ2 and PDZ3 interact with aPKC Kinase Domain-PBM (KD-PBM) module (19). Given that Par-3 PDZ2 is required to displace Cdc42 from the Par complex, we examined whether the aPKC PBM was also required for this activity. We found that removing the aPKC PBM (Par-6/aPKC ΔPBM) prevented Par-3 from displacing Cdc42 from the Par complex (Figure 3B). Our results indicate that Par-3 PDZ2 and aPKC PBM are necessary for Par-3 to disrupt the Cdc42–Par complex interaction (Figure 3C).

### Par-3 BR-PDZ2 and PDZ3 displace Cdc42 from the Par complex

Given the requirement of the aPKC KD-PBM for Par-3’s ability to displace Cdc42 from the Par complex, we examined whether the Par-3 domains that bind the KD-PBM (PDZ2 and PDZ3) were each sufficient for this activity. We recently discovered a conserved basic region (BR) at the N-terminal end of Par-3 PDZ2 that increases PDZ2’s affinity for the Par complex, so we also examined the effect of Par-3 BR-PDZ2 (19). We found that PDZ2 alone was sufficient to displace Cdc42 from the Par complex but was not as effective as PDZ1-APM such that some Par complex remained bound to solid phase Cdc42 (Figure 4A). In contrast, BR-PDZ2 displaced Cdc42 to a similar extent as PDZ1-APM and resulted in little to no Par complex associated with solid phase Cdc42 (Figure 4A). We conclude that Par-3 BR-PDZ2 can sufficiently displace Cdc42 from the Par complex.

**Figure 4.**
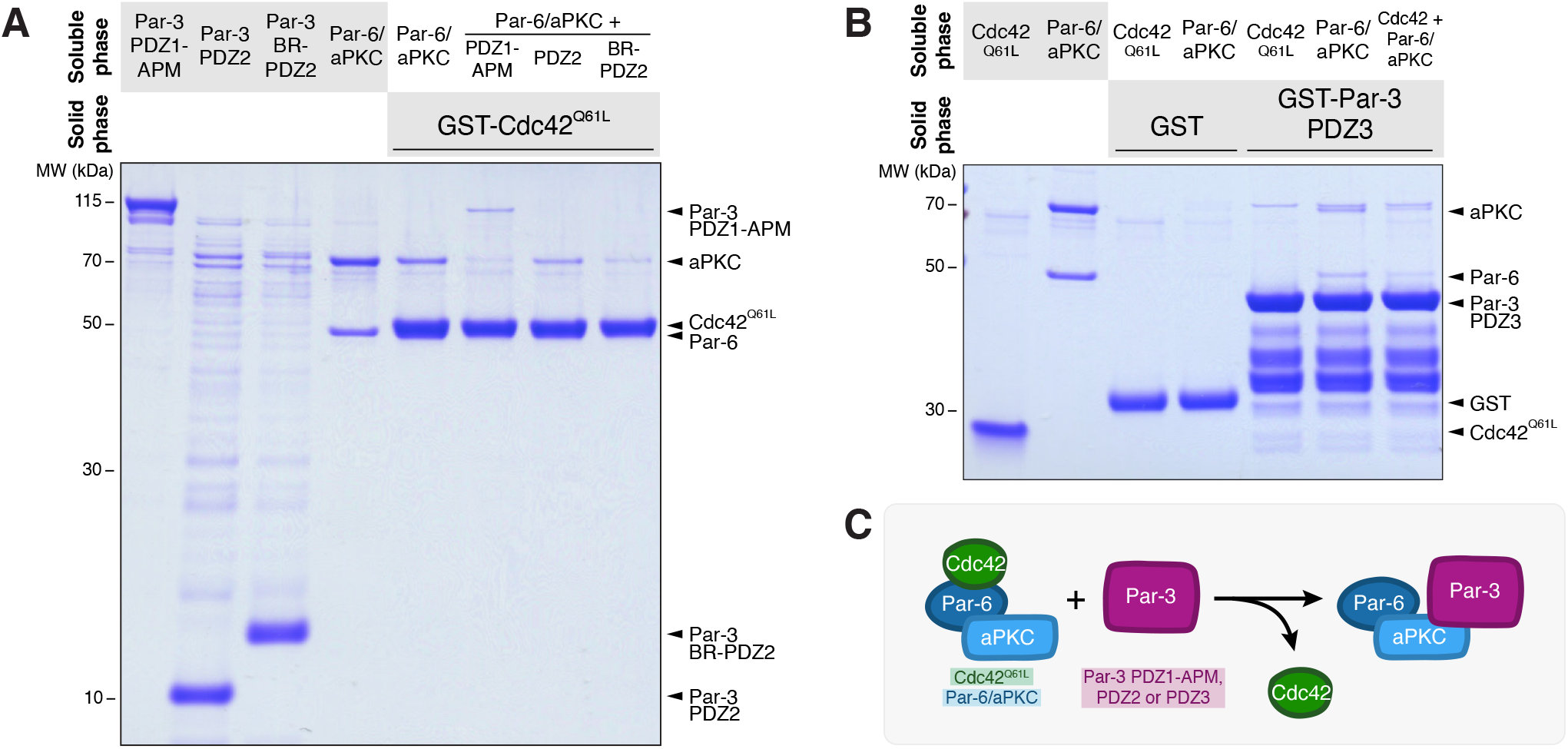
Par-3 PDZ2 and PDZ3 are sufficient for the displacement of Cdc42 from the Par complex. (**A**) Effect of Par-3 elements on the interaction between Cdc42 and the Par complex. Solid-phase (glutathione resin)-bound glutathione S-transferase (GST)-fused Cdc42^Q61L^ incubated with Par complex and/or Par-3 PDZ1-APM, PDZ2, or BR-PDZ2. Shaded regions indicate the fraction applied to the gel (soluble-phase or solid-phase components after mixing with soluble-phase components and washing). (**B**) Effect of Cdc42 on the interaction between Par-3 PDZ3 and the Par complex. GST-fused Par-3 PDZ3 incubated with Par complex and/or Cdc42^Q61L^. Labeling as described in (A). (**C**) Summary of Cdc42, Par-3, Par-6/aPKC elements that are sufficient for negative cooperativity between Par-3 and Cdc42 for the Par complex.

Like Par-3 PDZ2, Par-3 PDZ3 was found to interact with the Par complex utilizing a similar binding mode. Thus, we also tested the ability of Par-3 PDZ3 to displace Cdc42 from the Par complex. Given that Par-3 PDZ3 has a weak binding affinity for the Par complex (K_d_ of 78.9 μM) (19), we were unable to detect any significant change in the amount of Par complex bound to solid phase Cdc42 (data not shown). Therefore, we instead formed a Par-3 PDZ3-bound Par complex utilizing GST-Par-3 PDZ3 on the solid phase and soluble Par complex and examined the effect of Cdc42^Q61L^. We observed that addition of Cdc42 resulted in a reduction in the amount of Par complex bound to solid phase Par-3 PDZ3 indicating that Cdc42 is sufficient to displace PDZ3 from the Par complex (Figure 4B). Thus, our finding that Par-3 PDZ1-APM ΔPDZ2 cannot displace Cdc42 (Figure 3A) likely arises from the low binding affinity of PDZ3 for the Par complex compared to BR-PDZ2. Altogether, our results indicate that the Par-3 BR-PDZ2 and PDZ3—aPKC kinase-PBM (predominantly through BR-PDZ2) interactions negatively cooperate with the Cdc42—Par-6 CRIB-PDZ interaction (Figure 4C and 5A).

**Figure 5.**
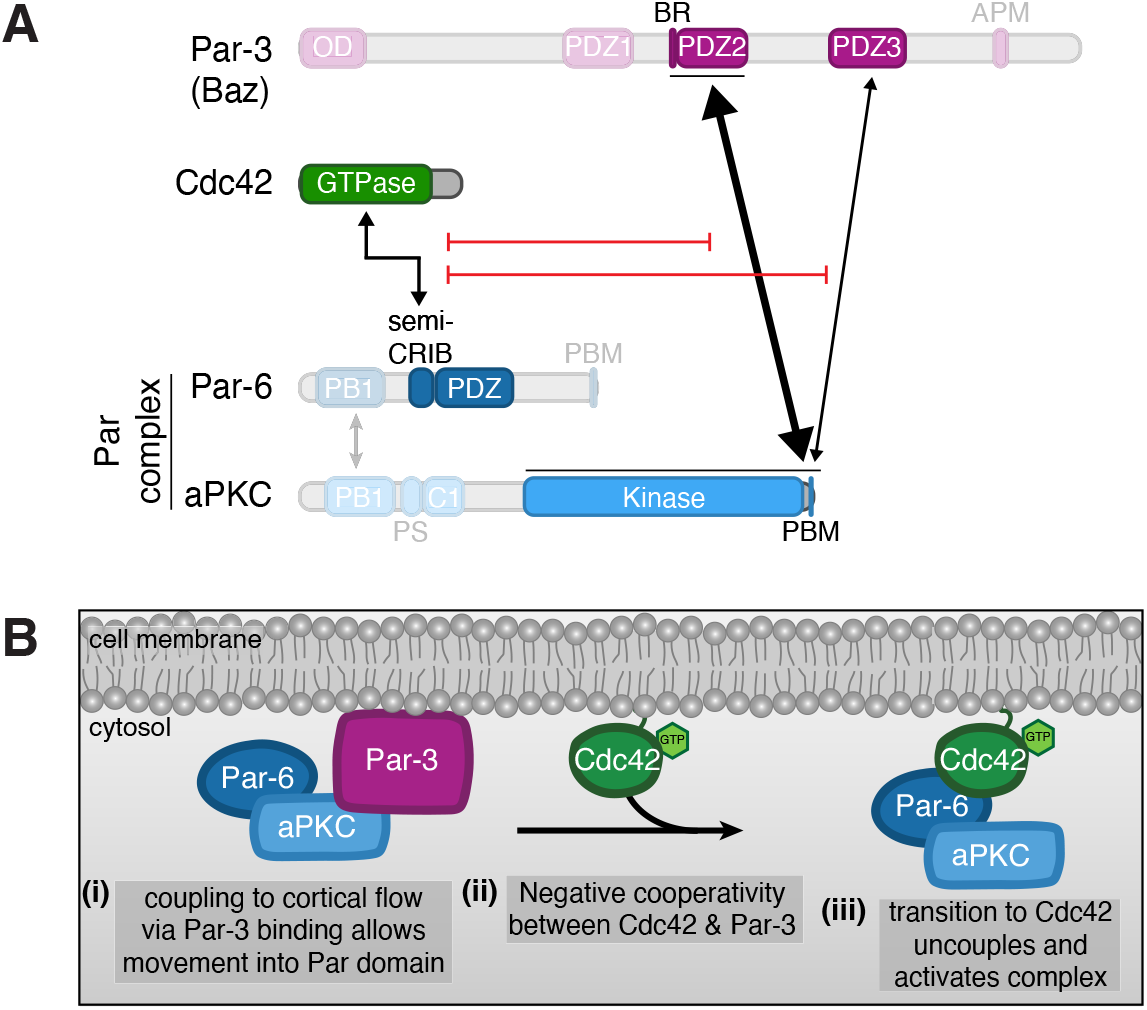
Model for transition from Par-3- to Cdc42-bound Par complex. (**A**) Domain architecture demonstrating the interactions between Par-3, Cdc42, and the Par complex that are sufficient for regulating the transition between a Par-3- and Cdc42-bound Par complex. (**B**) Model for polarization of the Par complex through the formation of distinct Par-3- and Cdc42-bound complexes. (i) Par-3-bound Par complex is coupled to cortical flow, allowing it to move towards the Par domain, (ii) active Cdc42 binds to Par-6 and inhibits the Par-3 BR-PDZ2/PDZ3—aPKC kinase-PBM interaction, resulting in the displacement of Par-3 from the Par complex and the formation of a Cdc42-bound Par complex, (iii) Cdc42-bound Par complex is polarized and active.

## Discussion

We examined how the Par complex transitions from Par-3- to Cdc42-bound states, a step that is thought to be critical for forming the Par cortical domain. In this model, the large size of oligomerized Par-3 couples the Par complex to actomyosin-driven directional cortical flows that transport the complex to its active domain (5, 20, 21). Once in this membrane region, the complex dissociates from Par-3 and binds Cdc42, allowing it to remain on the cortex and become activated. While the transition from Par-3 to Cdc42 is central to this model, little has been known about how switching between binding of each regulator occurs. We used a biochemical reconstitution approach to understand the mechanism of regulator switching, with the goal of identifying the minimal set of components required for the transition. A previous study that used co-immunoprecipation from cultured cell extracts concluded that Cdc42 and Par-3 can bind simultaneously to the Par complex, suggesting that other cellular components are required (9). For example, mechanical separation of the complexes by actomyosin-generated contraction has been proposed as one possibility (5, 20, 21). Using purified components, we found that external factors are not required as Cdc42 and Par-3 bind to the Par complex with strong negative cooperativity (Figure 5B). In this section, we examine the implications of our findings and speculate on key outstanding issues that remain in understanding this critical step in Par-mediated polarity.

How can the strong negative cooperativity between Cdc42 and Par-3 binding to the Par complex that we observed be reconciled with the previous observation of a quaternary complex? There are several possible explanations. First, it does not appear that ATP was included in the previous binding experiment, perhaps because it was not clear that Par-3 is an aPKC substrate at the time. We have found that the phosphorylation site on Par-3 can form a stalled complex with the aPKC kinase domain when ATP is not present (18, 26). Alternately, since the experiment was performed in extracts, it’s possible that additional factors were present that inhibit negative cooperativity, allowing Cdc42 and Par-3 to bind simultaneously. Finally, the presence of negative cooperativity does not necessarily preclude formation of a quaternary complex, albeit at reduced levels.

The nature of the Par complex interaction with Par-3 has been enigmatic because many distinct interactions have been reported (9–19). We examined which Par-3 interactions are coupled to Cdc42 binding and found that only one involving the aPKC kinase domain and PDZ binding motif (KD-PBM) is affected by Cdc42. This site binds with highest affinity to Par-3’s BR-PDZ2 domain but with weaker affinity to PDZ3 (19). Our results are consistent with the reported relative affinities – BR-PDZ2 most effectively reduces Cdc42 binding to the Par complex. We found that binding to the aPKC KD-PBM is both necessary and sufficient to displace Cdc42, and that Cdc42 can nearly completely displace Par-3 PDZ1-APM from the Par complex. These results indicate that the other reported interactions of Par-3 with the Par complex are not likely to be relevant to this step of Par-mediated cell polarity.

A central consequence of our findings for Par complex function is that the transition from Par-3-bound to Cdc42-bound Par complex does not require additional factors. Thus, the presence of Cdc42 alone could be sufficient for the transition to take place. However, our results do not preclude the possibility that other factors assist in complex switching. As we found that the affinity of Cdc42 for the Par complex is lower than that of Par-3, a higher concentration of Cdc42 would be required to achieve the same amount of Cdc42-bound complex. It is possible that the amount of Cdc42-bound complex does not need to be in excess of Par-3-bound complex, or that the concentration of active Cdc42 is higher than Par-3. Alternately, other factors could influence the affinity of Cdc42 with the Par complex – ligands of the Par-6 PDZ domain have been found to be coupled to Cdc42 binding, for example (28, 29).

How might the Cdc42 and Par-3 binding sites be coupled? The strongest negative cooperativity arises from a steric mechanism, where the binding sites would require steric overlap for Cdc42 and Par-3 to bind simultaneously. Alternately, in an allosteric mechanism binding is coupled to changes in structure or dynamics that reduce the affinity for the other regulator. While the binding site for Cdc42 is on Par-6 (semi-CRIB) and Par-3’s binding site is on aPKC (KD-PBM), little is known about the structural arrangement of the domains within the Par complex. Thus, it is formally possible that the semi-CRIB and KD-PBM are near one another, and it has been speculated that the PDZ domain adjacent to the semi-CRIB interacts with the KD-PBM (30). However, it is also clear that the Par complex is highly allosteric, as aPKC is autoinhibited from an intramolecular interaction between its pseudosubstrate and the kinase domain, and that Par-6 partially disrupts this interaction (31).

## Materials and Methods

### Protein Expression

#### Bacterial cells

Plasmids were transformed into BL21-DE3 cells, aliquoted onto LB + AMP plates, and grown at 37°C for 18 hours. Colonies were picked to inoculate 100 mL LB + AMP starter cultures and grown at 37°C for 2-3 hours until an OD_600_ of 0.4-1.0 was reached. Starter cultures were then diluted into 2 L LB + AMP, grown at 37°C to an OD_600_ of 0.8-1.0, and induced with 500 μM IPTG for 3 hours. Cultures were centrifuged at 5,000 RPM for 15-20 minutes and pellets were resuspended in nickel lysis buffer (50 mM NaH_3_PO_4_, 300 mM NaCl, 10 mM Imidazole, pH 8.0), GST lysis buffer (IX PBS, 1 mM DTT, pH 7.5), or maltose lysis buffer (20 mM Tris, 200 mM NaCl, 1 mM EDTA, 1 mM DTT, pH 7.5). Resuspended pellets were then frozen in liquid N_2_ and stored at −80°C.

#### Mammalian Cells

Par-6 and aPKC plasmids were co-expressed in FreeStyle 293-F cells as previously described (18, 19, 31). Briefly, cells were grown in FreeStyle 293 expression media in shaker flasks at 37°C with 8% CO_2_ and transfected with either 293fectin or ExpiFectamine (see Manufacturer’s protocol for more details). After 48 hours, cells were centrifuged 1-2x at 500 g for 3 minutes and cell pellets were resuspended in nickel lysis buffer (50 mM NaH_3_PO_4_, 300 mM NaCl, 10 mM Imidazole, pH 8.0). Resuspended pellets were then frozen in liquid N_2_ and stored at −80°C.

### Protein Purification

#### Bacterial cells

Resuspended pellets were thawed and then lysed by probe sonication at 70% amplitude, 0.3 seconds/0.7 seconds on/off pulse rate, 3x 1 minute. To pellet cellular debris, lysates were centrifuged at 15,000 RPM for 20 minutes. Lysates for GST-Cdc42^Q61L^ and GST-Par-3 PDZ3 were aliquoted, frozen in liquid N_2_ and stored at −80°C. For all other proteins, lysates were incubated with resin (amylose for most MBP-fused proteins and cobalt or nickel resin for His-fused proteins; MBP-Par-3 PDZ1-APM-His was His-purified) for 30-60 minutes at 4°C with mixing. Protein-bound resin was washed 3x with nickel lysis buffer (50 mM NaH_3_PO_4_, 300 mM NaCl, 10 mM Imidazole, pH 8.0) or maltose lysis buffer (20 mM Tris, 200 mM NaCl, 1 mM EDTA, 1 mM DTT, pH 7.5) and then eluted in 0.5-1.8 mL fractions with nickel elution buffer (50 mM NH_3_PO_4_, 300 mM NaCl, 300 mM Imidazole, pH 8.0) or maltose elution buffer (maltose lysis buffer, 10 mM Maltose). Protein-containing fractions were pooled and buffer exchanged into 20 mM HEPES, pH 7.5, 100 mM NaCl, and 1 mM DTT using a PD10 desalting column. Protein was then concentrated using a Vivaspin 20 centrifugal concentrator, aliquoted, frozen in liquid N_2_, and stored at −80°C.

#### Mammalian cells

Resuspended 293F pellets were thawed, lysed by probe sonication at 70% amplitude, 0.3 seconds/0.7 seconds on/off pulse rate, 4x 1 minute, and centrifuged at 15,000 RPM for 20 minutes. Lysates were incubated with resin (cobalt or nickel resin) for 30-60 minutes at 4°C with mixing. Protein-bound resin was washed 3x with nickel lysis buffer (50 mM NaH_3_PO_4_, 300 mM NaCl, 10 mM Imidazole, pH 8.0), with the first and second washes containing 100 μM ATP and 5 mM MgCl_2_. Protein was eluted in 0.5-0.6 mL fractions with nickel elution buffer (50 mM NH_3_PO_4_, 300 mM NaCl, 300 mM Imidazole, pH 8.0) and protein-containing fractions were then pooled together. Protein was buffer exchanged into 20 mM Tris, pH 7.5, 100 mM NaCl, 1 mM DTT, 100 μM ATP, and 5 mM MgCl_2_ using a PD10 desalting column and then purified by anion exchange chromatography using an AKTA FPLC protein purification system. Protein was filtered, injected into a Source Q column, and eluted with a salt gradient of 100-500 mM NaCl. Par complex-containing fractions were pooled and buffer exchanged into 20 mM HEPES, pH 7.5, 100 mM NaCl, 1 mM DTT, 100 μM ATP, and 5 mM MgCl_2_ using a PD10 desalting column. Protein was then concentrated using a Vivaspin 20 centrifugal concentrator, aliquoted, frozen in liquid N_2_, and stored at −80°C.

### Qualitative Binding Assay (Affinity Chromatography)

Bacterial lysates were incubated with resin (glutathione or amylose) for 30 minutes at 4°C and washed 4x with binding buffer (20 mM HEPES pH 7.5, 100 mM NaCl, 5 mM MgCl_2_, 0.5% Tween-20, 1 mM DTT, and 200 μM ATP). Soluble proteins were added to protein-labeled resin and incubated at room temperature with rotational mixing (incubation times of 10 minutes for MBP-Par-3 PDZ1-APM and 60 minutes for GST-fused proteins). Resin was washed 2-3x with binding buffer and proteins were eluted with 4X LDS sample buffer. Samples were run on a 12% Bis-Tris gel and stained with Coomassie Brilliant Blue R-250.

### Quantitative Binding Assay (Supernatant Depletion)

Bacterial lysates were incubated with resin (glutathione or amylose) for 30 minutes at 4°C and washed 6x (3x quick washes, 3x 5 minutes washes) with binding buffer (20 mM HEPES pH 7.5, 100 mM NaCl, 5 mM MgCl_2_, 0.5% Tween-20, 1 mM DTT, and 200 μM ATP). Two-fold serial dilutions of protein-bound resin were prepared with unlabeled resin as previously described (18, 19). Soluble protein (“receptor” or R) was added to protein-bound resin (“ligand” or L) and incubated at room temperature with rotational mixing (incubation times of 10 minutes for MBP-Par-3 PDZ1-APM and 60 minutes for GST-Cdc42^Q61L^). Samples containing only unlabeled resin and soluble protein were used as a negative control for binding. After incubation, samples were centrifuged, and an aliquot of the supernatant was collected and diluted in 4X LDS sample buffer. Samples were then run on a 12% Bis-Tris gel and stained with Coomassie Brilliant Blue R-250. Solid phase protein concentration was verified using a standard curve generated with known concentrations of a protein standard. All band intensities were quantified using ImageJ (v1.53a). The fraction of soluble phase protein (R) bound to solid phase protein (L) at a specific concentration of L ([L] = x) was determined with the following equation:

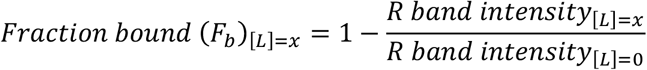

We then determined the dilution at which L resulted in 30-60% depletion (F_b_ = 0.3-0.6) and repeated the assay at this dilution in sextuplicate. Using F_b_, we determined the binding equilibrium dissociation constant (K_d_) with the following equation:

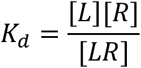

where [L] and [R] are the concentrations of free L and R at equilibrium and [LR] is the concentration of L bound to R.

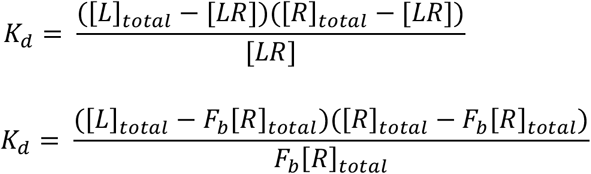

We also used the equation for the standard Gibbs free energy exchange to determine the binding energy of these interactions:

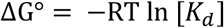

All data was analyzed using Microsoft Excel (v16.53), GraphPad Prism (v9.2), and DABEST (32).

## Key Resources Table

| Reagent or Resource type | Reagent or Resource | Source | Identifier | Additional Information |
| --- | --- | --- | --- | --- |
| Recombinant Protein | MBP-Par-3 PDZ1-APM | PMID: 32084408 |  | Expressed in BL21 cells from pMAL Par-3 309-987-His; His-purified |
| Recombinant Protein | MBP-Par-3 PDZ1-APM ΔPDZ1 | This paper | | Cloned by Q5 site-directed mutagenesis; expressed in BL21 cells from pMAL Par-3 393-987-His; His-purified |
| Recombinant Protein | MBP-Par-3 PDZ1-APM ΔPDZ2 | PMID: 35787373 | | Expressed in BL21 cells from pMAL Par-3 309-987 Δ437-533-His; His-purified |
| Recombinant Protein | MBP-Par-3 PDZ1-APM ΔPDZ3 | This paper | | Cloned by Q5 site-directed mutagenesis; expressed in BL21 cells from pMAL Par-3 309-987 Δ616-741-His; His-purified |
| Recombinant Protein | His-Par-3 PDZ2 | PMID: 35787373 |  | Expressed in BL21 cells from pET19 Par-3 444-533; His-purified |
| Recombinant Protein | His-Par-3 BR-PDZ2 | PMID: 35787373 |  | Expressed in BL21 cells from pET19 Par-3 426-533; His-purified |
| Recombinant Protein | GST-Par-3 PDZ3 | PMID: 35787373 |  | Expressed in BL21 cells from pGEX Par-3 616-741 |
| Recombinant Protein | His-Cdc42<sup>Q61L</sup> | This paper |  | Cloned by traditional methods; expressed in BL21 cells from pBH Cdc42 1-191 Q61L; His-purified |
| Recombinant Protein | GST-Cdc42<sup>Q61L</sup> | This paper |  | Cloned by traditional methods; expressed in BL21 cells from pGEX Cdc42 1-191 Q61L |
| Recombinant Protein | Par complex | PMID: 32084408 |  | Expressed in 293F cells from pCMV His Par-6 1-351 and pCMV aPKC 1-606; His-purified |
| Recombinant Protein | Par complex ΔaPKC PBM | PMID: 32084408 | | Expressed in 293F cells from pCMV His Par-6 1-351 and pCMV aPKC 1-600; His-purified |
| Recombinant DNA | pCMV (mammalian expression plasmid) | ThermoFisher | 10586014 |  |
| Recombinant DNA | pMAL c4X (bacterial expression plasmid) | Addgene | 75288 |  |
| Recombinant DNA | pGEX 4T1 (bacterial expression plasmid) | Amersham | 27458001 |  |
| Recombinant DNA | pBH (bacterial expression plasmid) | PMID: 15023337 |  |  |
| Recombinant DNA | pET19<br>(bacterial expression plasmid) | Millipore Sigma<br>(Novagen) | 69677 |  |
| Bacterial Strain | TG1 |  |  | Molecular cloning |
| Bacterial Strain | BL21-DE3 |  |  | Protein expression |
| Cell Line | FreeStyle 293-F | ThermoFisher | R79007 |  |
| Chemical | 293fectin Transfection Reagent | ThermoFisher | 12347019 |  |
| Chemical | ExpiFectamine 293 Transfection Kit | ThermoFisher | A14524 |  |
| Chemical | Freestyle 293 Expression Medium | ThermoFisher | 12338018 |  |
| Chemical | Expi293 Expression Medium | ThermoFisher | A1435101 |  |
| Chemical | Opti-MEM | ThermoFisher | 3198588 |  |
| Chemical | HisPur Cobalt Resin | ThermoFisher | 89965 |  |
| Chemical | HisPur NiNTA Resin | ThermoFisher | 88222 |  |
| Chemical | Amylose Resin | NEB | E8021L |  |
| Chemical | Glutathione Resin | GoldBio | G250-100 |  |
| Chemical | Source 30Q Anion Exchange Resin | GE Healthcare | 17-1275-01 |  |
| Chemical | 4X BOLT LDS Sample Buffer | ThermoFisher | B0007 |  |
| Chemical | PageRuler Plus Prestained Protein Ladder, 10-250 kDa | ThermoFisher | 26619 |  |
| Chemical | 20X BOLT MES SDS Running Buffer | ThermoFisher | B0002 |  |
| Chemical | Coomassie Brilliant Blue R-250 | GoldBio | C-461-5 |  |
| Chemical | Q5 Site-Directed Mutagenesis Kit | NEB | E0552S |  |
| Chemical | Gibson Assembly Cloning Kit | NEB | E5510S |  |
| Other | 125 mL Erlenmeyer Flasks | VWR | 89095-258 |  |
| Other | 250 mL Erlenmeyer Flasks | VWR | 89095-266 |  |
| Other | Bolt 12% Bis-Tris Gels | ThermoFisher | NW00125BOX |  |
| Other | PD-10 Desalting Columns | VWR | 95017-001 |  |
| Other | VivaSpin 20 MWCO 5kDa | Cytiva | 28932359 |  |
| Other | VivaSpin 20 MWCO 10kDa | Cytiva | 28932360 |  |
| Other | VivaSpin 20 MWCO 30kDa | Cytiva | 28932361 |  |
| Software | ImageJ | NIH |  | https://imagej.nih.gov/ij/ |
| Software | Prism | GraphPad Software |  | https://www.graphpad.com/ |
| Software | Estimation<br>Statistics BETA | PMID:<br>31217592 |  | www.estimationstats.com |

## Acknowledgments

This work was supported by NIH grant GM127092 (K.E.P.) and T32HD007348 (E.V.).

## References

1. Munro, E., Nance, J., and Priess, J. R. (2004) Cortical Flows Powered by Asymmetrical Contraction Transport PAR Proteins to Establish and Maintain Anterior-Posterior Polarity in the Early C. elegans Embryo. Dev. Cell. 7, 413–424

2. Hutterer, A., Betschinger, J., Petronczki, M., and Knoblich, J. A. (2004) Sequential Roles of Cdc42, Par-6, aPKC, and Lgl in the Establishment of Epithelial Polarity during Drosophila Embryogenesis. Dev. Cell. 6, 845–854

3. Aceto, D., Beers, M., and Kemphues, K. J. (2006) Interaction of PAR-6 with CDC-42 is required for maintenance but not establishment of PAR asymmetry in C. elegans. Dev. Biol. 299, 386–397

4. Atwood, S. X., Chabu, C., Penkert, R. R., Doe, C. Q., and Prehoda, K. E. (2007) Cdc42 acts downstream of Bazooka to regulate neuroblast polarity through Par-6 aPKC. J. Cell Sci. 120, 3200–3206

5. Rodriguez, J., Peglion, F., Martin, J., Hubatsch, L., Reich, J., Hirani, N., Gubieda, A. G., Roffey, J., Fernandes, A. R., St Johnston, D., Ahringer, J., and Goehring, N. W. (2017) aPKC Cycles between Functionally Distinct PAR Protein Assemblies to Drive Cell Polarity. Dev. Cell. 42, 400–415.e9

6. Oon, C. H., and Prehoda, K. E. (2019) Asymmetric recruitment and actin-dependent cortical flows drive the neuroblast polarity cycle. eLife. 8, e45815

7. Oon, C. H., and Prehoda, K. E. (2021) Phases of cortical actomyosin dynamics coupled to the neuroblast polarity cycle. eLife. 10, e66574

8. Lang, C. F., and Munro, E. (2017) The PAR proteins: from molecular circuits to dynamic self-stabilizing cell polarity. Development. 144, 3405–3416

9. Joberty, G., Petersen, C., Gao, L., and Macara, I. G. (2000) The cell-polarity protein Par6 links Par3 and atypical protein kinase C to Cdc42. Nat. Cell Biol. 2, 531–539

10. Lin, D., Edwards, A. S., Fawcett, J. P., Mbamalu, G., Scott, J. D., and Pawson, T. (2000) A mammalian PAR-3–PAR-6 complex implicated in Cdc42/Rac1 and aPKC signalling and cell polarity. Nat. Cell Biol. 2, 540–547

11. Qiu, R.-G., Abo, A., and Martin, G. S. (2000) A human homolog of the C. elegans polarity determinant Par-6 links Rac and Cdc42 to PKCζ signaling and cell transformation. Curr. Biol. 10, 697–707

12. Noda, Y., Takeya, R., Ohno, S., Naito, S., Ito, T., and Sumimoto, H. (2001) Human homologues of the *Caenorhabditis elegans* cell polarity protein PAR6 as an adaptor that links the small GTPases Rac and Cdc42 to atypical protein kinase C: Human PAR6s link Rac and Cdc42 to aPKC. Genes Cells. 6, 107–119

13. Garrard, S. M., Capaldo, C. T., Gao, L., Rosen, M. K., Macara, I. G., and Tomchick, D. R. (2003) Structure of Cdc42 in a complex with the GTPase-binding domain of the cell polarity protein, Par6. EMBO J. 22, 1125–1133

14. Izumi, Y., Hirose, T., Tamai, Y., Hirai, S., Nagashima, Y., Fujimoto, T., Tabuse, Y., Kemphues, K. J., and Ohno, S. (1998) An Atypical PKC Directly Associates and Colocalizes at the Epithelial Tight Junction with ASIP, a Mammalian Homologue of Caenorhabditis elegans Polarity Protein PAR-3. J. Cell Biol. 143, 95–106

15. Wodarz, A., Ramrath, A., Grimm, A., and Knust, E. (2000) Drosophila atypical protein kinase C associates with Bazooka and controls polarity of epithelia and neuroblasts. J. Cell Biol. 150, 1361–1374

16. Li, J., Kim, H., Aceto, D. G., Hung, J., Aono, S., and Kemphues, K. J. (2010) Binding to PKC-3, but not to PAR-3 or to a conventional PDZ domain ligand, is required for PAR-6 function in C. elegans. Dev. Biol. 340, 88–98

17. Renschler, F. A., Bruekner, S. R., Salomon, P. L., Mukherjee, A., Kullmann, L., Schütz-Stoffregen, M. C., Henzler, C., Pawson, T., Krahn, M. P., and Wiesner, S. (2018) Structural basis for the interaction between the cell polarity proteins Par3 and Par6. Sci. Signal. 11, eaam9899

18. Holly, R. W., Jones, K., and Prehoda, K. E. (2020) A Conserved PDZ-Binding Motif in aPKC Interacts with Par-3 and Mediates Cortical Polarity. Curr. Biol. 30, 893–898.e5

19. Penkert, R. R., Vargas, E., and Prehoda, K. E. (2022) Energetic determinants of animal cell polarity regulator Par-3 interaction with the Par complex. J. Biol. Chem. 298, 102223

20. Wang, S.-C., Low, T. Y. F., Nishimura, Y., Gole, L., Yu, W., and Motegi, F. (2017) Cortical forces and CDC-42 control clustering of PAR proteins for Caenorhabditis elegans embryonic polarization. Nat. Cell Biol. 19, 988–995

21. Dickinson, D. J., Schwager, F., Pintard, L., Gotta, M., and Goldstein, B. (2017) A Single-Cell Biochemistry Approach Reveals PAR Complex Dynamics during Cell Polarization. Dev. Cell. 42, 416–434.e11

22. Beers, M. (2006) Depletion of the co-chaperone CDC-37 reveals two modes of PAR-6 cortical association in C. elegans embryos. Development. 133, 3745–3754

23. Morais-de-Sá, E., Mirouse, V., and St Johnston, D. (2010) aPKC phosphorylation of Bazooka defines the apical/lateral border in Drosophila epithelial cells. Cell. 141, 509–523

24. Walther, R. F., and Pichaud, F. (2010) Crumbs/DaPKC-Dependent Apical Exclusion of Bazooka Promotes Photoreceptor Polarity Remodeling. Curr. Biol. 20, 1065–1074

25. Soriano, E. V., Ivanova, M. E., Fletcher, G., Riou, P., Knowles, P. P., Barnouin, K., Purkiss, A., Kostelecky, B., Saiu, P., Linch, M., Elbediwy, A., Kjær, S., O’Reilly, N., Snijders, A. P., Parker, P. J., Thompson, B. J., and McDonald, N. Q. (2016) aPKC Inhibition by Par3 CR3 Flanking Regions Controls Substrate Access and Underpins Apical-Junctional Polarization. Dev. Cell. 38, 384–398

26. Holly, R. W., and Prehoda, K. E. (2019) Phosphorylation of Par-3 by Atypical Protein Kinase C and Competition between Its Substrates. Dev. Cell. 49, 678–679

27. Pollard, T. D. (2010) A Guide to Simple and Informative Binding Assays. Mol. Biol. Cell. 21, 4061–4067

28. Peterson, F. C., Penkert, R. R., Volkman, B. F., and Prehoda, K. E. (2004) Cdc42 Regulates the Par-6 PDZ Domain through an Allosteric CRIB-PDZ Transition. Mol. Cell. 13, 665–676

29. Whitney, D. S., Peterson, F. C., Kittell, A. W., Egner, J. M., Prehoda, K. E., and Volkman, B. F. (2016) Binding of Crumbs to the Par-6 CRIB-PDZ Module Is Regulated by Cdc42. Biochemistry. 55, 1455–1461

30. Dong, W., Lu, J., Zhang, X., Wu, Y., Lettieri, K., Hammond, G. R., and Hong, Y. (2020) A polybasic domain in aPKC mediates Par6-dependent control of membrane targeting and kinase activity. J. Cell Biol. 219, e201903031

31. Graybill, C., Wee, B., Atwood, S. X., and Prehoda, K. E. (2012) Partitioning-defective Protein 6 (Par-6) Activates Atypical Protein Kinase C (aPKC) by Pseudosubstrate Displacement. J. Biol. Chem. 287, 21003–21011

32. Ho, J., Tumkaya, T., Aryal, S., Choi, H., and Claridge-Chang, A. (2019) Moving beyond P values: data analysis with estimation graphics. Nat. Methods. 16, 565–566

